# Modulation of serotonergic, dopaminergic and tubulin-associated pathways by tannin-free natural products in *Caenorhabditis elegans* reveals effective anthelmintic drug targets

**DOI:** 10.1101/2025.10.05.679051

**Authors:** Rajiur Rahaman Rabbi, Nurnabi Ahmed, Makshuder Rahman Zim, Mahfuzur Rahman Sajib, Mostak Ahmed, Babul Chandra Roy, Hasanuzzaman Talukder

## Abstract

Despite of the Southeast Asia’s rich medicinal flora, Bangladesh behinds in utilizing the *Caenorhabditis elegans* model for lead anthelmintic compound discovery. Moreover, target-specific resistance (TSR) and non-target-specific resistance (NTSR) surge have influenced the lead compounds bio-discovery over the past decades. To date, prior researches are limited to basic efficacy tests and conventional extraction protocol. Therefore, the study was designed to assess the anthelmintic potency of 19 tannin-free natural products (NPs) induced alterations of neurotransmission in *C. elegans*. The anthelmintic activity was assessed by examining the motility (head thrashes and body bends), mortality, egg hatch inhibition (EHI) and expression of *cat-1, ser-1, dat-1*, and *tba-1* genes. Eleven (11) plant extracts, *Azadirachta indica* A. Juss., *Cassia alata* L., *Portulaca oleracea* L., *Saraca asoca* (Roxb.) W.J.de Wilde, *Eleusine indica* L. Gaertn., *Persicaria hydropiper* L. Delarbre, *Foeniculum vulgare* Mill., *Clerodendrum infortunatum* L., *Linum usitatissimum* L., *Hibiscus rosa-sinensis* L., *and Vitex negundo* L. revealed a significant neurobehavioral and developmental impairments in *C. elegans. Persicaria hydropiper* L. Delarbre @ 1□mg/mL, caused the lowest body bending (31□±□1.7, *p*□<□0.01), while *Eleusine indica L. Gaertn*. reduced head thrashing (66.3□±□3.0, *p*□<□0.01). *L. usitatissimum* L. exhibited the highest lethality (87.3□±□1.8%, *p*□<□0.01) and the LD□□ values of *Eleusine indica* L. Gaertn. (0.40□mg/mL) and *Linum usitatissimum* L. (0.411□mg/mL) were the lowest. Interestingly, *C. infortunatum* L. exhibited the strongest EHI (97.5□±□3.0%, *p*□<□0.01). Gene expression analysis revealed a significant down-regulation of the dopaminergic and serotonergic pathway indicating the metabolic and reproductive disruptions. This pioneering study underscores the anthelmintic potency of native plant extracts, highlighting their neuromodulatory and developmental impact on *C. elegans*. Altogether, these findings suggest that the selected genes may serve as potential drug targets, paving the way for lead compounds identification and development to the advance anthelmintic drug discovery from the NPs.

## Introduction

Gastrointestinal nematodes (GINs) remain a major threat to livestock productivity and rural livelihoods in low- and middle-income countries (LMICs) such as Bangladesh [1]. These GI parasites impair nutrient absorption, cause chronic weight loss, reduce fertility which cause significant morbidity and even death if untreated [2]. Their impact is not limited to livestock but also infect more than estimated 3.5 billion people worldwide each year [3]. In livestock alone, the economic burden is substantial for example, intestinal worm infections in sheep alone cost the UK industry £83 million annually [4-7].

While synthetic anthelmintics remain the primary line of defense, decades of frequent, and often improper, usage have led to widespread anthelmintic resistance (AR) in both animal and human populations [8]. Recently, resistance has confirmed across all major drug classes including benzimidazoles, macrocyclic lactones, levamisole/imidazothiazoles, piperazine, paraherquamide, and amino-acetonitrile derivatives [9] and in various host species ranging from sheep and cattle to pigs and even humans [10]. One of the most striking cause of AR is widespread and error-prone dosing of these drugs threatening both human and animal health [11, 12]. However downstream mechanism of AR may arise through two mechanisms: target site resistance (TSR), where mutations alter drug-binding sites, and non-target site resistance (NTSR), which involves changes in drug metabolism, transport or elimination [13-15]. This resistance crisis severely threatens sustainable parasite control and food security in vulnerable regions.

One promising strategy for screening natural compounds is to target the nervous system, which is functionally distinct in nematodes compared to mammals. Nematode neurons are non-myelinated and reduced in number, making them vulnerable to neuroactive agents that can selectively disrupt parasite function without harming the host [16]. Neuro-synaptic genes such as *cat-1(*(involved in neurotransmitter vesicle transport), *ser-1* (a serotonin receptor), and *tba-1* (an alpha-tubulin essential for axonal integrity) play crucial roles in locomotion, feeding, and sensory response, making them valuable molecular markers for neurogenic anthelmintic action [17-19].

Several natural plants such as *Azadirachta indica A*.*Juss*.[20], *Calotropis procera* [21] and *Moringa oleifera* [22] have shown the promising anthelmintic activity. Based on the previous works on medicinal plants regarding phytochemical profiling (eg. HPLC, GC-MS etc.) of their active compounds with possible functional role [23-25] has been considered for in-vivo testing specifically for the antiparasitic activity. Despite of Bangladesh having a rich reservoir of medicinal flora, research on their antiparasitic potential using *Caenorhabditis elegans* as a model remains inconclusive [26]. Moreover, most of these studies remain limited to basic efficacy tests, lacking in-depth exploration of the molecular mechanisms involved. To investigate the anthelmintic potential of the selected medicinal plants from Bangladesh, we used *C. elegans* as a model nematode.

*C. elegans* is a cornerstone model organism that has made significant contributions to the fields of biology, including the developmental biology, neuroscience and cell biology. Increasingly, the scientific interest is to understanding its natural ecology and evolutionary lineage. It was the first multicellular organism to have its entire genome sequenced [27], and the first to have its complete neural wiring diagram or connectome mapped [28, 29]. This small, transparent nematode has a fixed number of somatic cells (959) and neurons (302), making it ideal for studies on cellular development, neurobiology, and behavior. The utility of *C. elegans* extends to a wide array of biological research. It has been effectively used to study the programmed cell death [30], muscle differentiation [31] and phytochemical screening via high-throughput screening [32, 33]. Its role in understanding the mode of action and resistance mechanisms of anthelmintic drugs [34-36], host–microbe interactions [37, 38], stress responses [39] toxicology [40, 41] and neurobiological disorders [42] further demonstrates its versatility. It is also proved to be a successful model in veterinary parasitology [43].

The present study was designed to screen the anthelmintic potency of natural products (NPs) from medicinal plants on *C. elegans* through phenotypic alterations and also to analyze the synaptic gene expression in *C. elegans* due to the treatment with NPs for understanding the possible mechanisms and their anthelmintic potency. By integrating behavioral phenotyping with gene expression profiling, this study aims to identify neurotropic, plant-derived compounds with therapeutic potential against resistant nematodes. Such an approach could contribute significantly to sustainable parasite control and natural drug discovery efforts in Bangladesh and beyond.

## Methods

### Plant material collection and identification

Plant materials were collected from the Botanical Garden, Germplasm Center of Bangladesh Agricultural University (BAU) campus and surrounding areas of BAU with the consent of the curators (S1 Fig). The plants were selected based on their ethno-pharmacological relevance and the corresponding information on each selected plant is presented in Table 1. The collection of plant material did not involve wild harvesting of threatened species on the IUCN Red List. All materials were obtained exclusively from cultivated plants maintained in the BAU Botanical Garden, authenticated by a botanist of the Department of Crop Botany, BAU and voucher specimens were deposited in the departmental herbarium. This study adheres to the IUCN Policy Statement on Research Involving Species at Risk of Extinction and does not pose any threat to natural populations of endangered species.

**Table 1.** Ethno-botanical data of selected medicinal plants having anthelmintic potency.

| **Id no** | **Local name of plants** | **Scientific name** | **Family** | **Reference** |
| --- | --- | --- | --- | --- |
| 1 | Neem(f) | *Azadirachta indica* A. Juss. | Meliaceae | [83] |
| 2 | Korola(f) | *Momordica charantia* L. | Cucurbitaceae | [84] |
| 3 | Kalosorisha(s) | *Mutarda nigra* Bernh. | Brassicaceae | [85] |
| 4 | Dad Mordon(l) | *Cassia alata* L. | Fabaceae | [86] |
| 5 | Korola(l) | *Momordica charantia* L. | Cucurbitaceae | [87] |
| 6 | Shialmuta(l) | *Ageratum conyzoides* L. | Asteraceae | [88] |
| 7 | Nunia(l) | *Portulaca oleracea* L. | Portulacaceae | [89] |
| 8 | Basok(l) | *Justicia adhatoda* L. | Acanthaceae | [90] |
| 9 | Asok(l) | *Saraca asoca* (Roxb.) W.J.de Wilde | Fabaceae | [91] |
| 10 | Chapra(l) | *Eleusine indica* L. Gaertn. | Poaceae | [92] |
| 11 | Bishkatali(l) | *Persicaria hydropiper* L. Delarbre | Polygonaceae | [91] |
| 12 | Chatim(l) | *Alstonia scholaris* L. R.Br. | Apocynaceae | [94] |
| 13 | Lebu(l) | *Citrus limon* L. Osbeck | Rutaceae | [95] |
| 14 | Mouri(s) | *Foeniculum vulgare* Mill. | Apiaceae | [96] |
| 15 | Vant(l) | *Clerodendrum infortunatum* L. | Lamiaceae | [97] |
| 16 | Tishi(s) | *Linum usitatissimum* L. | Linaceae | [98] |
| 17 | Methi(s) | *Trigonella foenum-graecum* L. | Fabaceae | [99] |
| 18 | Amloki(l) | *Phyllanthus emblica* L. | Phyllanthaceae | [100] |
| 19 | Joba(l) | *Hibiscus rosa-sinensis* L. | Malvaceae | [101] |
| 20 | Nishinda(l) | *Vitex negundo* L. | Lamiaceae | [22] |

Note: Here, (l)- leaf; (f)- fruit; s- seed

### Tannin-free extraction process of medicinal plants

Crude extracts using dichloromethane (CH□Cl□) and methanol (MeOH) were prepared on a small scale following earlier protocols [44, 45] with some alterations. About□300 mg of pulverized dried plant products was transferred to 15 mL falcon tube. For defatting, 5 mL n-hexane was added and mixed in a rotator for 1 minute, then centrifuged at 3000 rpm for 3 minutes and the supernatant was discarded while this process was subsequently repeated once. Next step, 7 mL dichloromethane (DCM) was added and mixed for 15 minutes under sonication (water bath with sonicator) at room temperature (RT). The liquid was filtered through Whatman filter-paper to collect the liquid in a fresh 15 mL falcon tube. To prepare the methanol (MeOH) fraction, 13 mL MeOH was added to the tube containing remaining materials. Again, mixed for 15 minutes under sonication at RT. The liquid was filtered again through Whatman filter-paper to collect the liquid in a fresh 15 mL falcon tube. Then an amount of 0.45g of PVPP (Polyvinylpolypyrrolidone) powder (Sigma-Aldrich, USA) was added to each tube. Then the tube was allowed to vortex for 10 s and put at 4°C for 15 min in a slow rotator. Then after 15 min, again it allowed to vortex for 10 s, then centrifuged at 3,000 g for 10 minutes. For each extract tube, the liquid + residue was filtered through Whatman filter-paper to collect the clear liquid in a fresh falcon tube. Then both tubes were mixed (Fig 1). Then the extract was sent to Professor Muhammed Hussain Central Laboratory (PMHCL) to evaporate the content to dryness using rotary evaporator and subsequently freeze drier. Then dry weight of extract was measured. DMSO (1%) was added to prepare the final concentration of 10 mg/mL.

**Fig 1.**
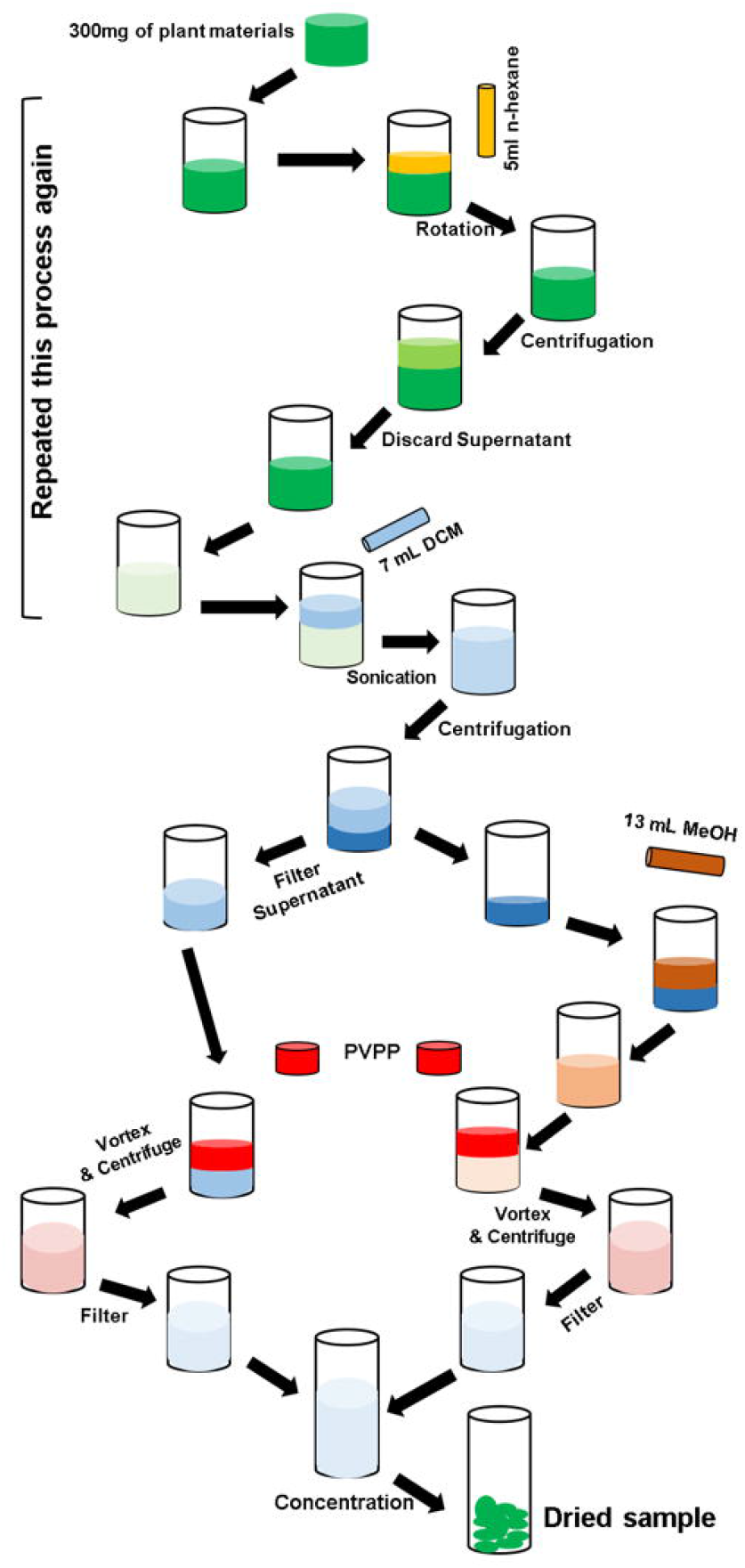
Schematic diagram of tannin-free extraction process of medicinal plants. PVPP: Polyvinylpolypyrrolidone

### Assay design

A balanced salt solution (M9) was used as the diluent for solvents and it was also used as the medium to prepare a nematode stock with 50 nematodes/20 µl. Synchronized population of *C. elegans* (N2) were collected using a standard bleaching assay and used in the phenotypic and genotypic screening. Tests were performed in 48-well plates (S2 Fig) containing a total volume of 250 µl/well, where nematodes were maintained on nematode growth medium (NGM) supplemented with *E. coli* OP50 as a food source. Each treatment was tested with five graded concentrations of plant extract (0.125–2 mg/ml) with three replicates per concentration. Levamisole (0.1 mg/ml) served as the positive control, while 1% DMSO was used as the negative control. Plates were incubated at 24 ºC for 24 h. Phenotypic screening assays were performed at least twice.

### *C. elegans* larvae exposure to extracts and motility assay

To evaluate the motility of treated *C. elegans*, the study measured head thrashes and body bends within a specific time frame. The methods were performed with modification as described previously [46]. To assay the head thrashes, *C. elegans* (N2) worms were synchronized to the L4 stage and then washed off from the cultivation plate. Every worm was transferred into a 48-well microtiter plate containing 60 µL of M9 buffer on the top of agar and the PE treatments. After a 1-min recovery in NGM containing PEs, the head thrashes were counted in a blinded manner for next 1 min. A thrash was defined as a change in the direction of bending at the mid-body. To evaluate body bend frequency, PEs treated worms were transferred to NGM plates, and the number of body bends was counted in a blinded manner over a 1-minute interval. A single body bend was characterized as a change in the direction of the posterior pharyngeal bulb along the y-axis, assuming the worm’s movement occurred along the x-axis. For each concentration of plant extract, observations were made manually on fifteen individual worms.

### Egg hatch inhibition assay (EHIA)

The *C. elegans* (N2) was cultured on nematode growth medium (NGM) plates seeded with *E. coli* (OP50). Worms were harvested by centrifugation at 1,000 rpm for 1 minute, followed by two washes with phosphate-buffered saline (PBS) and subsequently resuspended in fresh PBS. Approximately 0.5 mL of synchronized gravid adults was transferred into a 15 mL centrifuge tube containing 5 mL of hypochlorite solution and incubated for 5 minutes to dissolve adult worms, leaving only the eggs intact. The eggs were pelleted by centrifugation at 1,000 rpm for 1 minute, washed thrice with PBS, and resuspended in 10 mL PBS. Approximately 40–60 eggs were aliquoted into each well of a 48-well plate containing 100 µL PBS with plant extracts at a concentration of 1 mg/mL. Control wells contained eggs suspended in PBS without extracts. Plates were incubated at 24 °C for 48 hr. Post-incubation, Lugol’s iodine solution was added and the wells were examined under an inverted microscope to enumerate both eggs and hatched larvae [47]. Egg hatch inhibition (%) was calculated based on the formula given below.

Percent inhibition (%) = 100 (1 −*P*_test_/*P*_non-treated_), Where *P* is the number of eggs that hatched in EHIA.

### Mortality assay

The mortality assay was carried out in 48-well microtiter plates (NGM), each containing 250 µL of PEs at different concentrations, with three replicates per concentration. Controls included 1% DMSO in M9 buffer as a negative control and 0.1 mg/mL levamisole in M9 as a positive control. Initial screening involved testing extract concentrations ranging from 0.125 to 2 mg/mL on *C. elegans* to identify doses causing significant lethality. The concentration showing the most substantial nematocidal effect was selected for detailed study, based on the approach described by previous study [48]. Each well was seeded with approximately 40–50 L4-stage nematodes and incubated at 24 °C. Anthelmintic activity was evaluated after 12, 24, and 48 hr of exposure to the extracts. Worms were classified as dead if they remained straight, translucent and exhibited no movement of the pharynx, head or tail for at least 10 seconds [49]. The median lethal concentration (LC□□) for most effective and also for Levamisole (0.025-0.125 mg/mL) was then determined, with viability assessments primarily conducted at 24 hr.

### Synaptic gene expression analysis in *C. elegans* after treatment with NPs

The neuromodulator genes *ser-1, cat-1, tba-1*, and *dat-1* were selected for analysis based on prior studies and publicly available microarray expression data [18, 19] with *actin-1* used as the reference (housekeeping) gene. L4 or newly adult-stage *C. elegans* were exposed to 0.5□mg/mL of plant extracts for 24 hr. Following incubation, worms were washed three times with NGM buffer to eliminate residual bacteria. Actively moving worms were then transferred into microcentrifuge tubes and subsequently total RNA was extracted according to the manufacturer’s protocol of Favor Prep™ Total RNA Isolation Kit (Biotech Corp. Taiwan). The concentration of RNA was determined using the Multiskan SkyHigh Microplate Spectrophotometer (Thermo Fisher Scientific Inc. USA). Total RNA (<1 μg) was used for the synthesis of cDNA in 20 μL reaction by ProtoScript^®^ II First Strand cDNA Synthesis Kit (New England Biolabs. USA). The quantitative expression of six gene transcripts was examined by qRT-PCR. The qRT-PCR assays were carried out in a 20 μL reaction mixture containing 5 μL template cDNA, 0.5 μL (10 μM/μL) of both forward and reverse primers, 10 μL 2×Luna^®^ Universal qPCR Master Mix (New England Biolabs, USA) and 4 μL RNase-free water. The reaction was started with the initial activation of the polymerase at 95°C for 1 min, followed by denaturation of the template at 95°C for 15s; annealing and elongation were done at 60ºC for 30 s. After the completion of 45 cycles, the melting curve was achieved at 60 to 95ºC to assess the presence of a unique final product. The primers used in RT-qPCR were listed in Table 2. Using the comparative threshold cycle (ΔΔCT) approach, the mRNA concentration of the interested gene was measured in relation to *actin-1* expression.

**Table 2.**
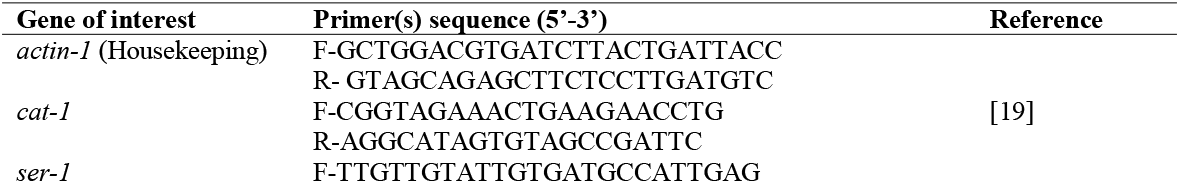
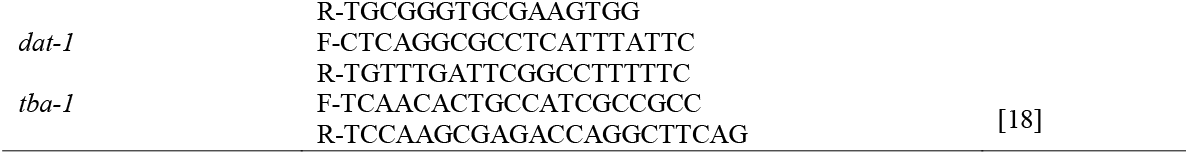
Primer(s) used for qPCR amplification of targeted genes.

## Results

### Natural products (NPs) significantly reduce the motility of *C. elegans*

Significant reductions in head thrashing of *C. elegans* in response to various plant extracts at a concentration of 1 mg/mL compare to 1% DMSO was observed for *Azadirachta indica A*.*Juss*., *Cassia alata L*., *Ageratum conyzoides L*., *Saraca asoca (Roxb*.*) W*.*J*.*de Wilde, Eleusine indica L. Gaertn*., *Persicaria hydropiper L. Delarbre, Foeniculum vulgare, Clerodendrum infortunatum L*., *Linum usitatissimum L*., *Hibiscus rosa-sinensis L*., *Vitex negundo L*. (*p* < 0.01) and Levamisole (positive control) indicating a strong inhibitory effect on motility (Fig 2A). In contrast, extracts such as *Momordica charantia* L., *Mutarda nigra Bernh*., *Portulaca oleracea L*., *Justicia adhatoda L*., *Alstonia scholaris L. R*.*Br*., *Citrus limon L. Osbeck, Trigonella foenum-graecum L*.and *Phyllanthus emblica L*.showed non-significant effects, suggesting these extracts do not have significant effect on motility. The positive control, also showed a reduced head thrashing count, consistent with its anthelmintic action, while the overall data highlights the potential of certain plant extracts to influence *C. elegans* behavior.

**Fig 2.**
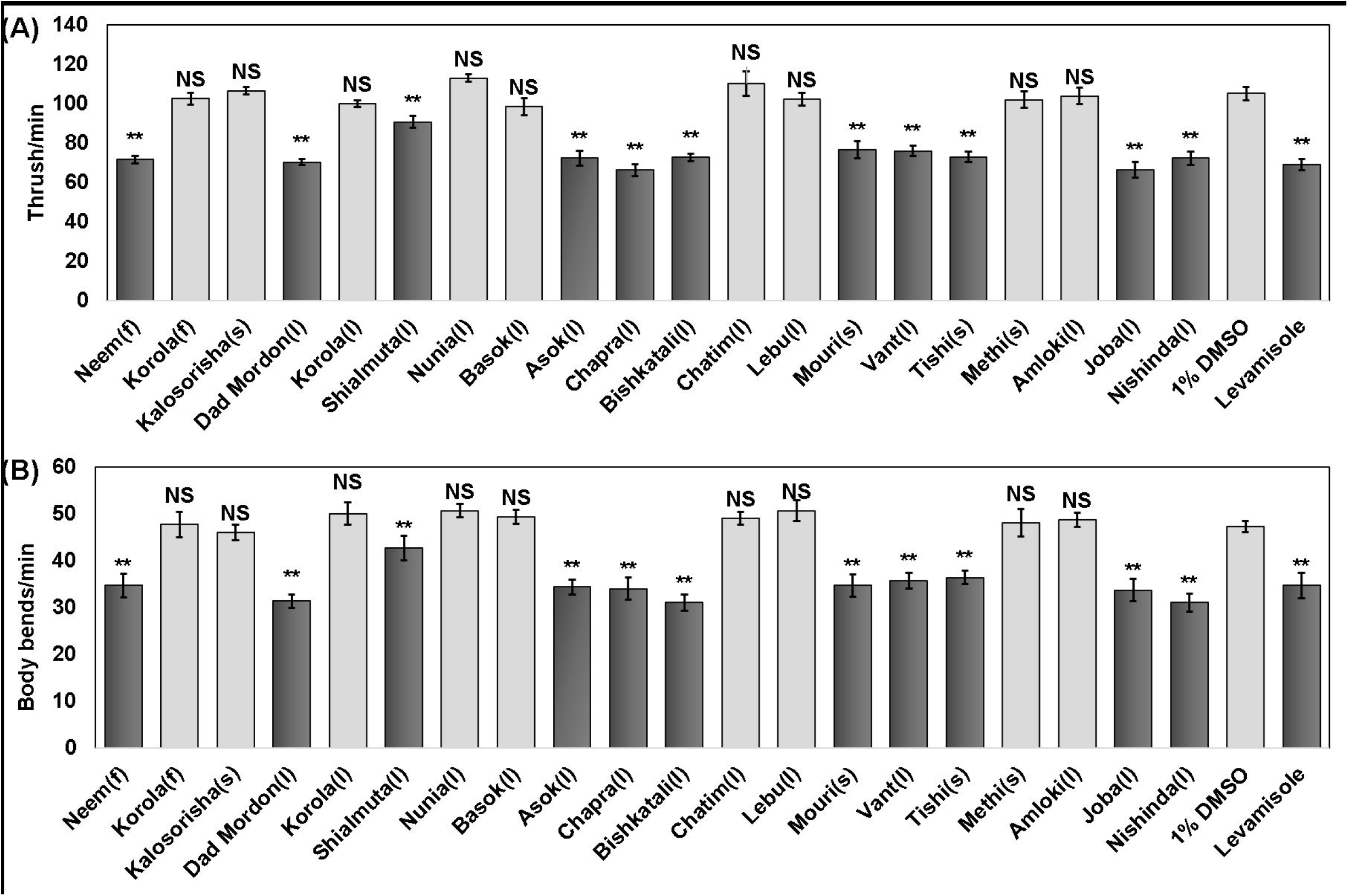
Effect of natural products on *C. elegans* motility. (A) Head thrush and (B) Body bends per minute. Data within the bar diagram presented as mean±SEM. Statistical significance compared the means between each concentration’s negative control. l, Leaves; r, roots; s, seeds, NS, not-significant. ^**^ Significantly different from negative control (1% DMSO) at P < 0.01.

Similarly, extracts including *Azadirachta indica A*.*Juss*., *Cassia alata L*., *Ageratum conyzoides L*., *Saraca asoca (Roxb*.*) W*.*J*.*de Wilde, Foeniculum vulgare, Clerodendrum infortunatum L*., *Linum usitatissimum L*., *Hibiscus rosa-sinensis L*., *Vitex negundo L*. and the positive control (Levamisole), exhibited significantly reduced body bending (p < 0.01), suggesting a strong inhibitory effect on the motility (Fig 2B). Additionally, *Eleusine indica L. Gaertn*. and *Persicaria hydropiper L. Delarbre* showed moderate reductions (p < 0.05). In contrast, other extracts such as *Momordica charantia* L., *Mutarda nigra Bernh*., *Portulaca oleracea L*., *Justicia adhatoda L*., *Alstonia scholaris L. R*.*Br*., *Citrus limon L. Osbeck, Trigonella foenum-graecum L*.and *Phyllanthus emblica L*.did not exhibit significant reductions in body bending, indicating minimal impact on *C. elegans* motility.

### Potent nematocidal activity of selected NPs

Variable biological responses were observed with the plant extracts at 1□mg/mL after 24hr. Worms in Tishi (*Linum usitatissimum L*.) exhibited the highest mortality rate at 87.33±1.76%, followed by Joba (*Hibiscus rosa-sinensis L*.) with 83.33±2.40% and Chapra (*Eleusine indica L. Gaertn*.) at 83.00±3.61%. These mortality rates were significantly higher than the negative control (1% DMSO, 1.00±0.0%). On the other hand, Korola (*Momordica charantia L*.) and Kalosorisha (*Mutarda nigra Bernh*.) exhibited relatively lower mortality (33.00±2% and 9.00±1%) respectively. However, Korola and Kalosorisha still demonstrated statistically significant effects compared to the negative control. Other extracts such as korola and Shialmuta (*Ageratum conyzoides L*.) also showed moderate mortality rates (49.50±0.5% and 51.50±1.5%) respectively. These findings highlight the varying levels of efficacy among the plant extracts, such as *Azadirachta indica A*.*Juss*., *Cassia alata L*., *Ageratum conyzoides L*., *Saraca asoca (Roxb*.*) W*.*J*.*de Wilde, Eleusine indica L. Gaertn*., *Persicaria hydropiper L. Delarbre, Foeniculum vulgare, Clerodendrum infortunatum L*., *Linum usitatissimum L*., *Hibiscus rosa-sinensis L*. and *Vitex negundo L*. showing strong nematocidal activity on *C. elegans*, indicating their potentiality as natural anthelmintic (Table 3).

**Table 3.** Mortality rate and egg hatch inhibition (%EHI) of selected PEs (1 mg/mL) on *C. elegans* at 24 h.

| Plant Name (local) | Mortality% | %EHI |
| --- | --- | --- |
| Neem(f) | 82.34±2.027*** | 93.75±0.0*** |
| Korola(f) | 33.00±2*** | 43.75±6.25*** |
| Kalosorisha(s) | 9.00±1 | 9.375±3.12 |
| Dad Mordon(l) | 79.50±3.97*** | 96.875±3.12*** |
| Korola(l) | 49.50±0.5*** | 18.75±6.25 |
| Shialmuta(l) | 51.50±1.5*** | 43.75±6.25*** |
| Nunia(l) | 82.67±3.53*** | 84.375±3.12*** |
| Basok(l) | 8.50±1.5 | 37.5±6.25*** |
| Asok(l) | 79.00±4.62*** | 93.75±0.0*** |
| Chapra(l) | 83.00±3.61*** | 90.625±3.12*** |
| Bishkatali(l) | 79.33±5.49*** | 93.75±0.0*** |
| Chatim(l) | 8.50±1.5 | 34.375±3.12*** |
| Lebu(l) | 7.00±2 | 31.25±6.25*** |
| Mouri(s) | 82.00±2.52*** | 90.625±3.12*** |
| Vant(l) | 77.33±5.46*** | 97.5±3.0*** |
| Tishi(s) | 87.33±1.76*** | 90.625±3.12*** |
| Methi(s) | 8.50±1.5 | 68.125±3.12*** |
| Amloki(l) | 22.50±2.5*** | 68.75±6.25*** |
| Joba(l) | 83.33±2.40*** | 84.375±3.12*** |
| Nishinda(l) | 82.00±3.51*** | 90.625±3.12*** |
| Lev 0.1mg/mL | 61.50±1.5*** | 79.125±9.37*** |
| 1% DMSO | 1.00±0.0 | 6.25±0.0 |

Note: Data presented as means ± SEM and are compared to 1% DMSO as the negative control. Statistical significance was calculated using a one-way ANOVA. Statistical significance compared the means between each concentration's negative
control and treatment. l, Leaves; r, roots; s, seeds; Lev, Levamisole. \*\*\* Significantly different from negative control (1% DMSO) at P < 0.001.

For these promising plants (11) and Levamisole (0.025-0.125 mg/mL), the LC_50_ values were calculated. The significant anthelmintic effects of various plant extracts were observed across a range of concentrations (0.125 to 2 mg/mL). At the highest concentration (2 mg/mL), the mortality rates for all plant extracts were substantial, with ranges 95 to 96%. Natural products of *Azadirachta indica A*.*Juss*., *Cassia alata L*., *Ageratum conyzoides L*., *Saraca asoca (Roxb*.*) W*.*J*.*de Wilde, Eleusine indica L. Gaertn*., *Persicaria hydropiper L. Delarbre, Foeniculum vulgare, Clerodendrum infortunatum L*., *Linum usitatissimum L*., *Hibiscus rosa-sinensis L*. and *Vitex negundo L*. all showed mortality rates in this range, with Vant (*Clerodendrum infortunatum L*.) achieving the highest at 96% (Table 4). At 1 mg/mL, mortality ranged from 79.5% to 83%, with similar trends observed at 0.5 mg/mL. Statistical analyses indicated that all plant extracts at these concentrations were significantly more effective than the negative control (1% DMSO), which showed only 1% mortality, confirming the efficacy of the plant natural compounds. The PEs show varying levels of efficacy, with Nishinda (*Eleusine indica L. Gaertn*.) having the lowest LC_50_ value of 0.40, indicating the highest potency among the tested plants. Tishi (*Linum usitatissimum L*.) Asok (*Saraca asoca Roxb. W*.*J*.*de Wilde*) and Mouri (*Foeniculum vulgare*) were also effective, with LC_50_ values of 0.411, 0.413, and 0.420 respectively, all grouped as significantly more effective than others. Plants like *Azadirachta indica A*.*Juss*. (Neem), *Cassia alata L*. (Dadmordon), *Ageratum conyzoides L*. (Nunia) and *Clerodendrum infortunatum L*. (Vant) have intermediate LC_50_ values around 0.43, suggesting the moderate anthelmintic activity. On the other hand, Bishkatali (*Persicaria hydropiper L. Delarbre*) and Joba (*Hibiscus rosa-sinensis L*.) exhibited the highest LC_50_ values of 0.492 and 0.459, indicating comparatively inferior efficacy relative to other PEs. However, when it comes for the figure of Levamisole, it was 0.058 indicating lowest LC_50_ value among the tested (Table 4).

**Table 4.** Dose-dependent mortality and LC_50_values of selected plant extracts on *C. elegans*.

| Plant name<br>(local) | Dose(mg/mL) |  |  |  |  | LC<sub>50</sub> |
| --- | --- | --- | --- | --- | --- | --- |
|  | 0.125 | 0.25 | 0.5 | 1 | 2 |  |
| Neem (f) | 35.67±4.05*** | 39.34±2.96*** | 55.34±2.34*** | 82.34±2.027*** | 95.5±2.18*** | 0.421 |
| Dadmordon (l) | 35.33±2.60*** | 40.00±3.21*** | 56.67±2.85*** | 79.50±3.97*** | 95.00±2.31*** | 0.423 |
| Nunia (l) | 35.00±5.03*** | 40.00±0.58*** | 54.00±2.65*** | 82.67±3.53*** | 93.00±1.53*** | 0.424 |
| Asok (l) | 30.00±3.21*** | 45.33±2.40*** | 58.67±3.76*** | 79.00±4.62*** | 95.17±0.93*** | 0.413 |
| Chapra (l) | 35.67±2.91*** | 42.00±1.73*** | 55.00±1.53*** | 83.00±3.61*** | 94.00±2.08*** | 0.40 |
| Bishkatali (l) | 33.33±1.76*** | 40.33±2.96*** | 49.67±1.45*** | 79.33±5.49*** | 93.33±1.86*** | 0.492 |
| Mouri (s) | 35.33±3.48*** | 40.67±2.03*** | 54.67±1.45*** | 82.00±2.52*** | 93.00±1.15*** | 0.420 |
| Vant (l) | 33.00±2.65*** | 45.00±2.08*** | 54.33±3.38*** | 77.33±5.46*** | 96.00±1.15*** | 0.429 |
| Tishi (s) | 31.00±1.53*** | 40.33±2.60*** | 58.00±1.00*** | 87.33±1.76*** | 93.67±0.88*** | 0.411 |
| Joba (l) | 30.67±2.73*** | 41.33±1.86*** | 53.67±1.86*** | 83.33±2.40*** | 92.33±1.76*** | 0.458 |
| Nishinda (l) | 33.67±1.76*** | 43.67±1.76*** | 56.67±2.33*** | 82.00±3.51*** | 95.67±1.20*** | 0.459 |
| Lev(0.1mg/mL) | 61.50±1.5*** |  |  |  |  | 0.058 |
| 1% DMSO | 1.0±0.001 |  |  |  |  |  |

Note: Data are presented as means ± SEM and are compared to 1% DMSO as the negative control. Statistical significance compared the means between each concentration's negative control and treatment. It was calculated using a one-way ANOVA. To calculate the LC<sub>50</sub> of Levamisole, graded concentrations (0.025-0.125mg/mL) were used. l, Leaves; r, roots; s, seeds; Lev, Levamisole. \*\*\* Significantly different from negative control (1% DMSO) at P < 0.001.

### Egg hatch inhibition (EHI)

The EHI values further underscore the anthelmintic potential of the tested plant extracts. Dad Mordon (*Cassia alata L*.) and Bishkatali (*Persicaria hydropiper L. Delarbre*) showed the highest EHI, with Dad Mordon achieving 96.875±3.12% and Bishkatali showing 93.75±0.0%, both significantly higher than the negative control (1% DMSO) and almost similar to the positive control (Levamisole, 79.125±9.37%). Other extracts, including Asok (*Saraca asoca Roxb. W*.*J*.*de Wilde*), Vant (*Clerodendrum infortunatum L*.), and Chapra (*Eleusine indica L. Gaertn*.), also exhibited substantial EHI (around 90% to 93.75%), reinforcing the observation that these plant species have considerable effects on inhibiting egg hatch. On the lower end, Kalosorisha (*Mutarda nigra Bernh*.), Basok (*Justicia adhatoda L*.), and Methi (*Trigonella foenum-graecum*) demonstrated lower EHI values, but still showed a significant inhibition compared to the negative control, with values ranging from 31.25±6.25% to 68.125±3.12%. Interestingly, Methi showed a notable EHI of 68.125±3.12%, which suggests a moderate, but significant anthelminthic potency (Table 3).

### Alteration of synaptic transduction in *C. elegans* after treatment with NPs

The *tba-1*gene expression analysis revealed strong downregulation (0.1–0.2) in *Persicaria hydropiper L. Delarbre* and *Eleusine indica L*., moderate (0.3-0.5) in *Portulaca oleracea L*., and poor (>0.5) in *Cassia alata* (0.6116) but still significant (p<0.01). For *ser-1, Portulaca oleracea L*. and *Saraca asoca (Roxb*.*) W*.*J*.*de Wilde* showed strong downregulation, *Azadirachta indica A*.*Juss*. and *Cassia alata L*. remained within the strong range but closer to moderate. In *cat-1*, strong downregulation was found in *Saraca asoca (Roxb*.*) W*.*J*.*de Wilde* and *Eleusine indica L*., while others were not included due to higher expression. For *dat-1*, strong downregulation occurred in all six extracts, with the lowest in *Persicaria hydropiper L. Delarbre, Portulaca oleracea L*. and *Saraca asoca (Roxb*.*) W*.*J*.*de Wilde* and relatively higher yet still strong downregulation in *Cassia alata L*. and *Azadirachta indica A*.*Juss*.

To gain insight into the molecular mechanisms underlying neurotransmission changes, the mRNA expression of specific stress-associated genes was assessed based on the findings above. The worms were treated to plant extracts to the newly adult stage (L4), the mRNA levels of serotonin-related genes such as *cat-1*(Enables dopamine:sodium symporter activity and serotonin:sodium symporter activity), *ser-1*(Enables G protein-coupled serotonin receptor activity and serotonin binding activity) and *tba-1* (Encodes tubulin, a key component of the cytoskeleton, essential for cellular structure and function) decreased significantly versus the non-exposed group. Similarly, the mRNA level dopamine-related gene *dat-1* decreased significantly (Fig 3B). The overall changes in gene expressions caused by extracts exposure could immediately reveal the stress response *in vivo*. In conjunction with the physiological consequences, the suppression of neurotransmitter production was primarily responsible for the stress responses exerted by the extracts of selective natural products. Furthermore, the mRNA level of *cat-1* and *dat-1* decreased drastically, and the expression of other gene was also reduced significantly (Fig 3D). These findings demonstrated further that the aberrant expression of the serotonin-, dopamine-, and microtubules formation-related genes likely contributed to the behavior, cellular, metabolic and reproductive impairments of *C. elegans* due to exposure to plant extracts (Fig 4).

**Fig 3.**
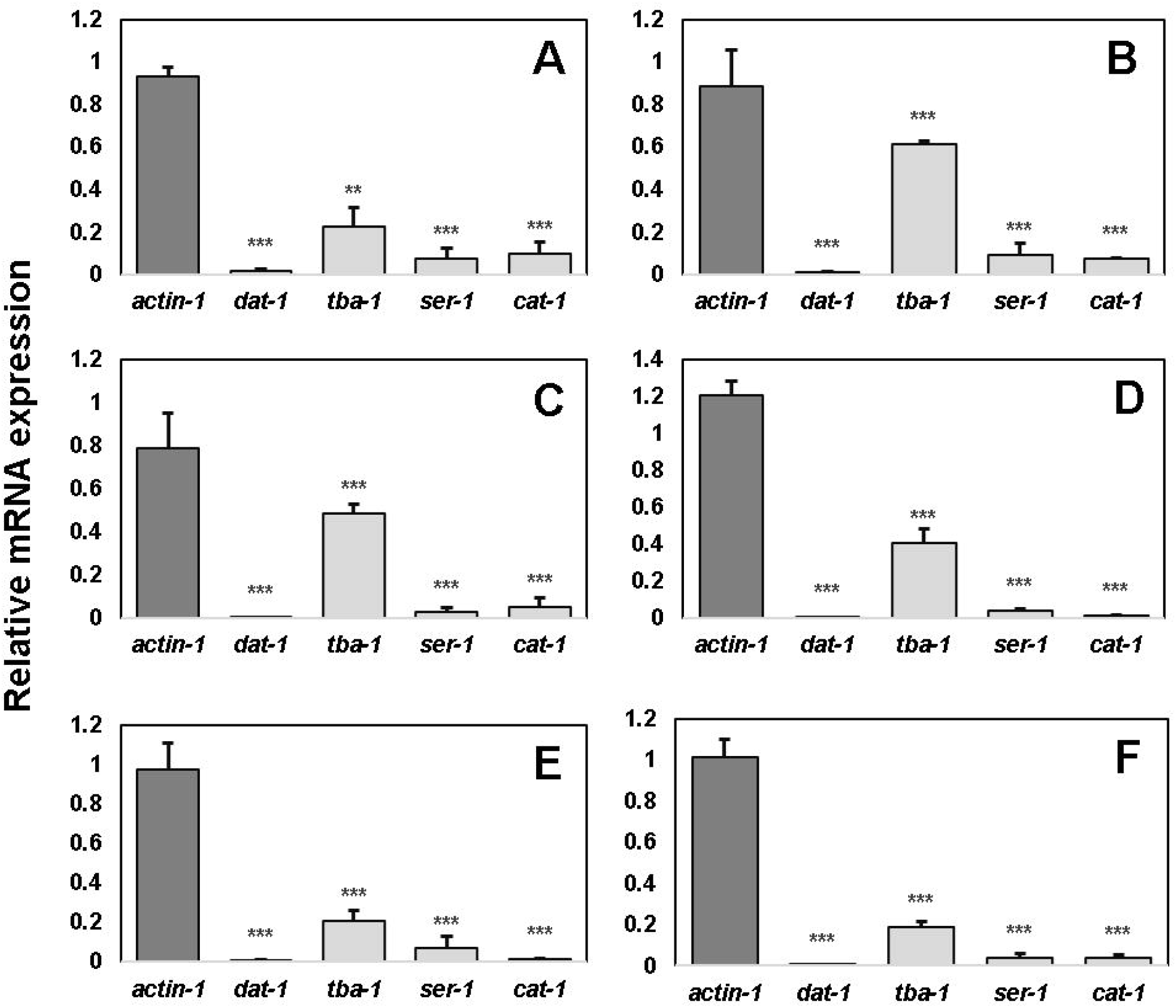
Analysis of relative mRNA expression of neurotransmission-related genes in *C. elegans* exposed to plant extracts. Exposure to six extracts decreased mRNA levels of selected genes. (A) Serotonin, dopamine and microtubule-related genes including *cat-1, ser-1, dat-1* and *tba-1* expression in response to Neem (*Azadirachta indica A. Juss*); (B) Dadmordon (*Cassia alata L*.); (C) Nunia (*Portulaca oleracea L*.); (D) Asok (*Saraca asoca Roxb. W*.*J*.*de Wilde*); (E) Chapra (*Eleusine indica L. Gaertn*.); (F) Bishkatali (*Persicaria hydropiper L. Delarbre*) extract at 0.5mg/mL. Values of neurotoxicity-related genes expressions were normalized using actin-1 mRNA and represent means relative to the control. The values are presented as mean ± SEM. *p*-value indicates the difference with DMSO-treated control. ns means not significant, * *p <* 0.05, ** *p <* 0.01, *** *p <* 0.001, Plant extracts treatment vs. control.

**Fig 4.**
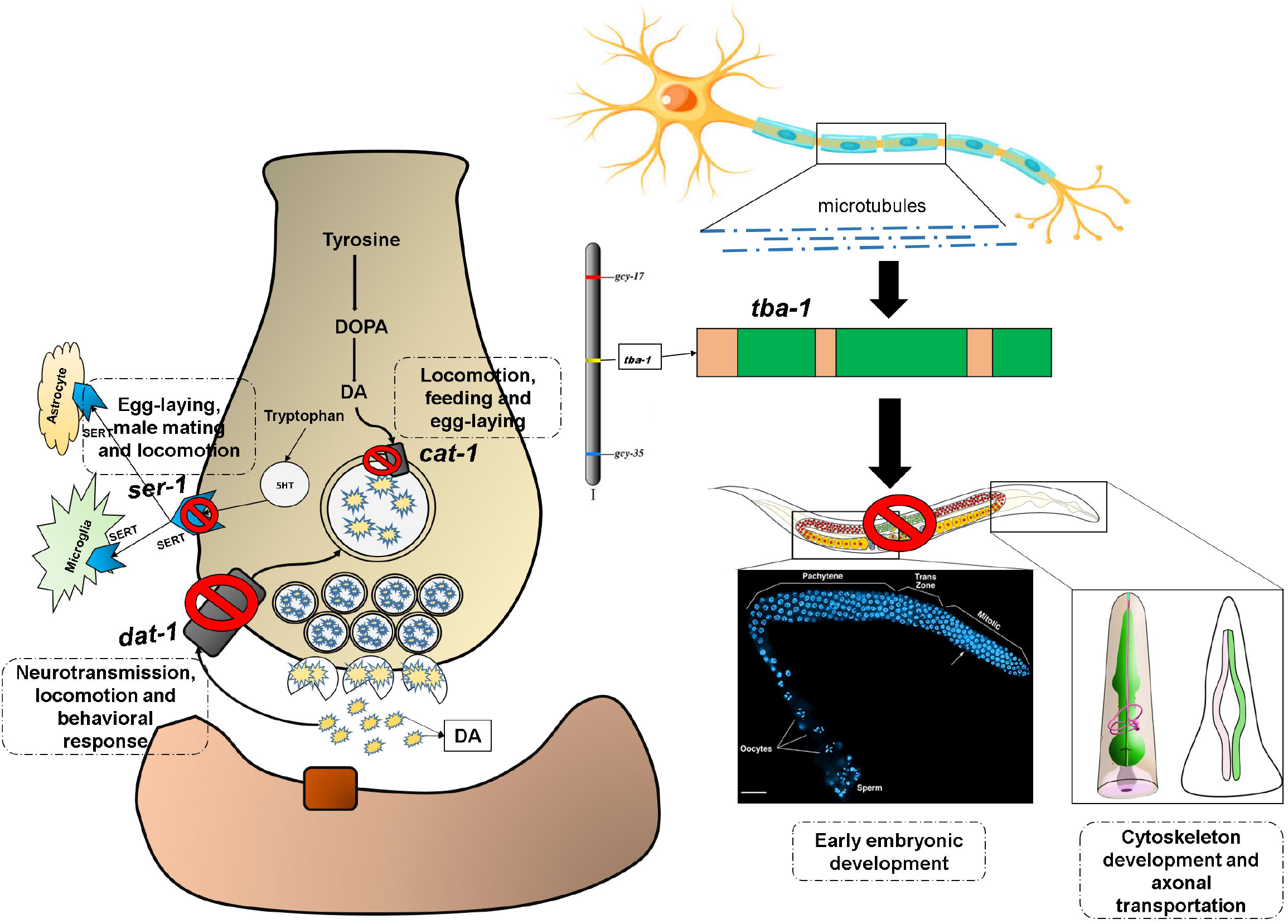
Proposed mechanism of action of natural products on *C. elegans*. The schematic illustration of neural transmission and Chromosome I of *C. elegans* highlighting known and predicted genes associated with neurophysiology, reproduction and body morphology. Here, three genes *cat-1, ser-1*, and *dat-1* were identified as being expressed in presynaptic neurons, where they play critical roles in regulating neurotransmitter signaling linked to locomotion and reproductive behaviors. Additionally, the *tba-1* gene, located both in axonal microtubules and chromosome I, is involved in early embryonic development, cytoskeletal organization and axonal transport. The tested natural products were found to interfere (marked with red blockage sign) with these molecular pathways, thereby altering the associated phenotypic traits.

## Discussion

Anthelmintic resistance (AR) has emerged as a global issue, primarily driven by two mechanisms, target-specific resistance (TSR) and non-target-specific resistance (NTSR). In the fact of this growing challenge, Bangladesh holds a unique advantage due to its rich biodiversity and abundance of medicinal plants, supported by favorable geographical conditions. This biodiversity offers valuable opportunities for the discovery of new therapeutics. To maximize the potential of NPs, this study is prioritized the selection of 19 plants based on their ethnopharmacological relevance. Plants with a history of traditional use are not only more likely to show biological activity [50, 51], but they are also generally considered safer and less toxic [52] compared to randomly selected plants. However, working with crude plant extracts can be challenging and often time-consuming. One major difficulty is the presence of bulk substances like lipids, waxes, chlorophyll and tannins, which often interfere with biological tests and make it harder to identify the active compounds. To address this, the study highlights the importance of using a carefully designed extraction process that removes these interfering substances and enriches the extract with potentially active ingredients. Additionally, the study developed a simple and efficient extraction method that works well even with limited amounts of plant material, making it ideal for rare or seasonal species. This practical approach provides a solid foundation for future research on plant-based anthelmintic agents.

Research aimed at discovering new anthelmintic drugs is costly, it has yet led to the identification of a considerable number of novel compounds over the past decade [53, 54]. The *C. elegans* has become a widely used model nematode for studying drugs against parasitic infections because it shares many genomic similarities with them [55]. Additionally, it offers an affordable and efficient system for drug screening. *C. elegans* model system is using globally for the suitability and advantages over parasitic nematodes. In this study, *C. elegans* model was used successfully for the first time in Bangladesh for screening the natural plant products to observe anthelmintic potency. The study identified eleven (11) plant extracts (*Azadirachta indica A*.*Juss*., *Cassia alata L*., *Portulaca oleracea L*., *Saraca asoca Roxb. W*.*J*.*de Wilde, Eleusine indica L. Gaertn*., *Persicaria hydropiper L. Delarbre, Foeniculum vulgare, Clerodendrum infortunatum L*., *Linum usitatissimum L*., *Hibiscus rosa-sinensis L*. and *Vitex negundo L*.) with significantly lower body bends and head thrushes at the tested concentration of 1mg/mL. Body bending and head thrash assays were employed in this study as indicators of motility. Significant reduction in response to the eleven extracts indicate the wide array chemical components or mixtures of those NPs leads to impair the locomotion. The present findings are in line with the previous studies where the reduction in body bends and head thrush indicative to disruption of neuro-behavioral function determined at indicative concentration [56-58].

The present study demonstrated that plant extracts exhibit varying levels of lethality on *C. elegans*, with several extracts showing significant anthelmintic activity. At a concentration of 1 mg/mL, *Linum usitatissimum L*. (Tishi) exhibited the highest mortality rate followed by *Hibiscus rosa-sinensis L*. (Joba) and these mortality rates suggest strong nematocidal potential. The LC_50_ values of the most potent extracts, such as *Eleusine indica L. Gaertn*. (Chapra), *Linum usitatissimum L*. (Tishi), *Azadirachta indica A*.*Juss*.(Neem) and *Cassia alata L*.(Dad Mordon), ranged between 0.40-0.43 mg/mL, indicating strong anthelmintic efficacy. These research align with previous research that highlights the nematocidal properties of plant-based bioactive compounds. Several studies have documented the anthelmintic potential of *Azadirachta indica A*.*Juss*.(Neem), which is rich in bioactive compounds like azadirachtin, nimbin and nimbidin known to disrupt nematode neuromuscular function and reproduction [23, 59, 60]. The potent activity observed in *Cassia alata L*. (Dad Mordon) is consistent with earlier findings that identified anthraquinones as the primary bioactive compounds responsible for nematocidal effects [61, 62]. The significantly high mortality rates observed at 2 mg/mL support the dose-dependent efficacy of plant extracts. Comparable results have been reported in studies on *Moringa oleifera* and *Allium sativum*, where increasing concentrations led to enhanced nematocidal activity due to oxidative stress and neurotoxicity in nematodes [63, 64]. The LC□□ values of the most effective extracts indicate higher potency than several previously studied medicinal plants, such as *Lespedeza cuneata, Salix X sepulcralis, Robinia pseudoacacia, Acer rubrum, Rosa multiflora, Quercus alba* and *Rhus typhina* which demonstrated LC□□ values exceeding 0.7mg/mL [65].

Interestingly, the moderate efficacy of *Hibiscus rosa-sinensis L*. (Joba) and *Persicaria hydropiper L. Delarbre* (Bishkatali) despite high mortality at 1 mg/mL suggests the presence of bioactive compounds that require longer exposure or specific molecular targets to exert maximum effects. This observation is consistent with reports on flavonoid-rich plants like *Citrus limon L. Osbeck* and *Foeniculum vulgare*, which show moderate LC□□ values but prolonged nematostatic effects [66, 67]. Notably, *Cassia alata L*. (Dad Mordon) and *Persicaria hydropiper L. Delarbre* (Bishkatali) showing the highest egg hatch inhibition (EHI), surpassing Levamisole at given concentration. Similar efficacy was observed in *Saraca asoca (Roxb*.*) W*.*J*.*de Wilde*(Asok), *Clerodendrum infortunatum L*. (Vant), and *Eleusine indica L. Gaertn*. (Chapra), indicating the potent anthelmintic activity. These findings are consistent with previous reports of *Cassia alata L*. Inhibiting nematode egg hatching due to their rich phytochemical composition, including flavonoids, tannins and alkaloids [68, 69]. Conversely, *Justicia adhatoda L*. (Basok), *Mutarda nigra Bernh. Bernh. Bernh*.(Kalosorisha), and *Trigonella foenum-graecum L*.(Methi) exhibited moderate EHI, suggesting a weaker but still significant effect. Studies indicate that their bioactive compounds, such as vasicine and saponins, contribute to nematocidal activity but at higher concentrations [70, 71].

The anthelmintic action of benzimidazoles (BZs) involves β-tubulin binding [72], which hinders microtubule formation, reduces glucose uptake and leads to energy exhaustion and parasite mortality. Levamisole acts as a nicotinic acetylcholine receptor (nAChR) agonist [73], causing continuous muscle contraction, paralysis, and expulsion of nematodes. The mode of action of macrocyclic lactones (e.g., ivermectin) involves activation of GluCl channels, resulting in elevated chloride ion influx, neuromuscular inhibition and eventual death [74]. While BZs starve parasites, levamisole paralyzes them, and MLs disrupt neural function, making these drugs effective against various helminths and arthropods. However, the growing reports of resistance to these anthelmintics are a significant concern [75]. This highlights the urgent need for new classes of anthelmintics with novel modes of action and also drug target to effectively combat parasitic infections. The study selected four genes which directly associated with neurotransmission as a potential drug targets. With this study, results on the expression of genes associated with neurotransmission, cytoskeletal integrity and stress response reveal the neurophysiological effects of six plant extracts (*Azadirachta indica A*.*Juss*., *Cassia alata L*., *Portulaca oleracea L*., *Saraca asoca (Roxb*.*) W*.*J*.*de Wilde, Eleusine indica L. Gaertn*., and *Persicaria hydropiper L. Delarbre*) in *C. elegans*. Notably, *tba-1*, which encodes tubulin, was down-regulated in worms treated with *Eleusine indica L. Gaertn*. and *Persicaria hydropiper L. Delarbre*, indicate potential disruption of microtubule stability, which could impair neuronal function and metabolism. These findings are aligned with the previous studies where it shown that plant-derived compounds can alter the cytoskeletal dynamics, leading to disrupted neuronal integrity and impaired motor function of the parasite [76]. Similarly, *ser-1*, a G-protein coupled serotonin receptor gene, showed the lowest expression in *Portulaca oleracea L*. and *Saraca asoca (Roxb*.*) W*.*J*.*de Wilde*, suggesting a possible decline in serotonergic signaling, which may affect behaviors like locomotion and feeding [77]. This result is consistent with studies indicating that plant extracts can modulate serotonergic pathways, either enhancing or inhibiting serotonin receptor function, influencing behaviors and mood [78-80]. The *cat-1* gene, involved in catecholamine biosynthesis, was suppressed in *Saraca asoca (Roxb*.*) W*.*J*.*de Wilde* and *Eleusine indica L. Gaertn*., pointing to potential disruptions in neurotransmitter synthesis. Previous research has shown that some plant compounds can interfere with neurotransmitter biosynthesis, which may have neurotoxic implications. Additionally, *dat-1*, which regulates dopamine reuptake, was significantly down-regulated in all plant extract-exposed groups, particularly in *Portulaca oleracea L*., *Saraca asoca (Roxb*.*) W*.*J*.*de Wilde*, and *Persicaria hydropiper L. Delarbre*, suggesting impaired dopaminergic signaling. This is consistent with studies where plant extracts led to the down-regulation of dopamine transporters, resulting the increased dopamine levels and potential alterations in motor and behavioral functions [81, 82]. These results collectively highlight the potential neurotoxic effects of certain plant extracts, particularly *Portulaca oleracea L*., *Saraca asoca W*.*J*.*de Wilde*, and *Persicaria hydropiper L. Delarbre*, which may disrupt key neurophysiological processes and supports the previous result of motility and mortality assay. Altogether, these findings extend the evidence from the present studies highlighting the importance of neurotransmission disruption in the modulation of neurobehavioral and developmental functions.

The emergence of resistance to existing anthelmintic drugs has also created an urgent need to explore novel drug targets that are essential, parasite-specific and less prone to resistance development. Identifying and validating such targets offers a promising path toward the design of next-generation therapeutics with enhanced efficacy and safety. Unlike conventional broad-spectrum approaches, novel anthelmintic targets enable precise disruption of critical biological pathways in helminths while minimizing harm to the host and environment. This paradigm shift is vital not only for sustaining parasite control in both human and veterinary medicine but also for safeguarding global food security and public health in the face of evolving parasitic threats. The present study, four validated genes, *ser-1, cat-1, tba-1*, and *dat-1* hypothesized as promising effective targets for anthelmintic drugs. These genes critically involvement in serotonergic and dopaminergic signaling pathways as well as cytoskeletal stability, showed strong and significant downregulation following treatment, correlating with marked impairment in *C. elegans* motility, viability and reproductive capacity. Their essential roles in neuromuscular coordination and survival, coupled with the observed phenotypic effects upon suppression, highlight their therapeutic relevance. The functional validation of these targets established a foundation for the development of targeted interventions and reinforces the potential of pathway-specific strategies in overcoming the growing threat of anthelmintic resistance. Thus, *ser-1, cat-1, tba-1*, and *dat-1* represent a new generation of validated, mechanism-informed drug targets for future anthelmintic discovery. This ground breaking study presents a robust and rapid method for NPs extraction, high-throughput screening and an effective approach to elucidate the molecular pathways underlying their mode of action, highlighting their potential as a reliable source for lead compound discovery. Given the large number of plant extracts screened (n = 19) with the predicted function of their NPs, comprehensive phytochemical profiling (e.g., HPLC, LC-MS) was not included. Instead of, the emphasis took place on functional and molecular assays to identify the promising candidates. Further studies should aim to optimize the high-throughput screening methods, discover further bioactive compounds and investigation of their therapeutic potentials in treating parasitic diseases. This study sets the stage for future advancements in parasitology, providing a solid foundation for the development of novel and sustainable treatments for parasitic infections.

## Conclusion

In this study, an optimized protocol of natural product extraction has been developed that can further enhance research on screening anthelmintic potency of NPs. Among the 19 selected plants, 11 species namely *Azadirachta indica A*.*Juss*., *Cassia alata L*., *Portulaca oleracea L*., *Saraca asoca (Roxb*.*) W*.*J*.*de Wilde, Eleusine indica L. Gaertn*., *Persicaria hydropiper L. Delarbre, Foeniculum vulgare, Clerodendrum infortunatum L*., *Linum usitatissimum L*., *Hibiscus rosa-sinensis L*., and *Vitex negundo L*. demonstrated notable anthelmintic efficacy. These plants significantly reduced body bending and head thrashing, indicating diminished motility and disruption of neuronal signal transduction. Additionally, the *C. elegans* exposure to NPs, increased mortality and inhibited egg hatching, suggesting potent nematocidal and ovicidal/larvicidal effects. The down regulation of *cat-1, ser-1, dat-1*, and *tba-1* genes in *C. elegans* further emphasizes the modulation of serotonergic and dopaminergic pathways, along with the suppression of metabolic and reproductive processes. These results support the selected genes can also be noble target of antiparasitic drugs. This study highlights the inaugural effort to conduct a high-throughput screening of NPs from selected medicinal plants, along with a detailed investigation of their mode of action on *C. elegans* in Bangladesh. However, in spite of challenges for funding, results of this study will help to identify and unveil lead compounds from these medicinal plants as well as their mechanisms in advancing novel anthelmintic discovery from natural products and further exploration of their potential in clinical applications.

## Supporting information

https://doi.org/10.5281/zenodo.17190978

## Acknowledgments

The authors gratefully acknowledge Dr. Muntasir Kamal, Department of Molecular Genetics, University of Toronto, Canada for generously providing the bacterial strains and N2 (wild type) *C. elegans*.

## Supporting information

**S1 Fig.**
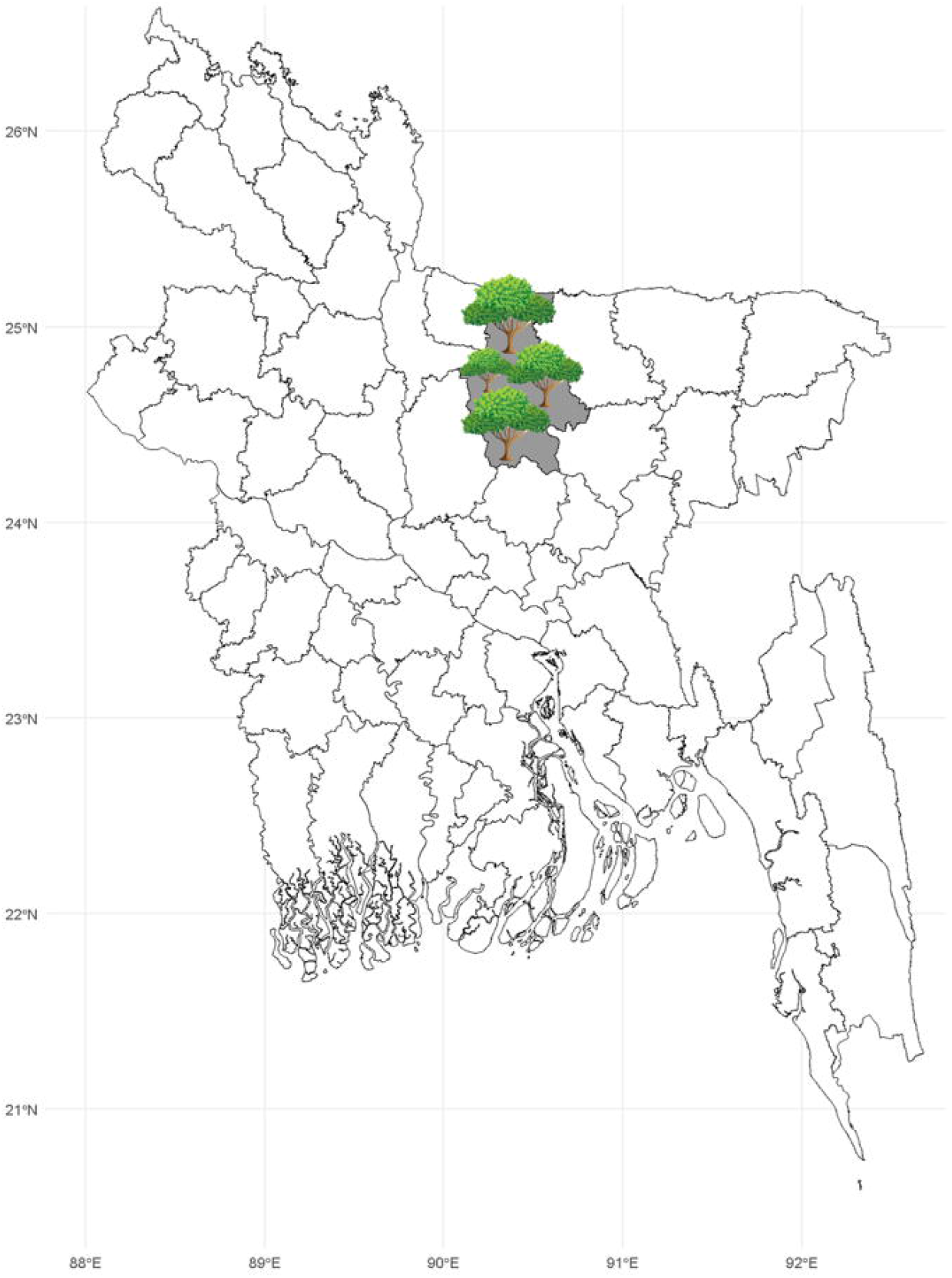
Medicinal plant collection area.

**S2 Fig.**
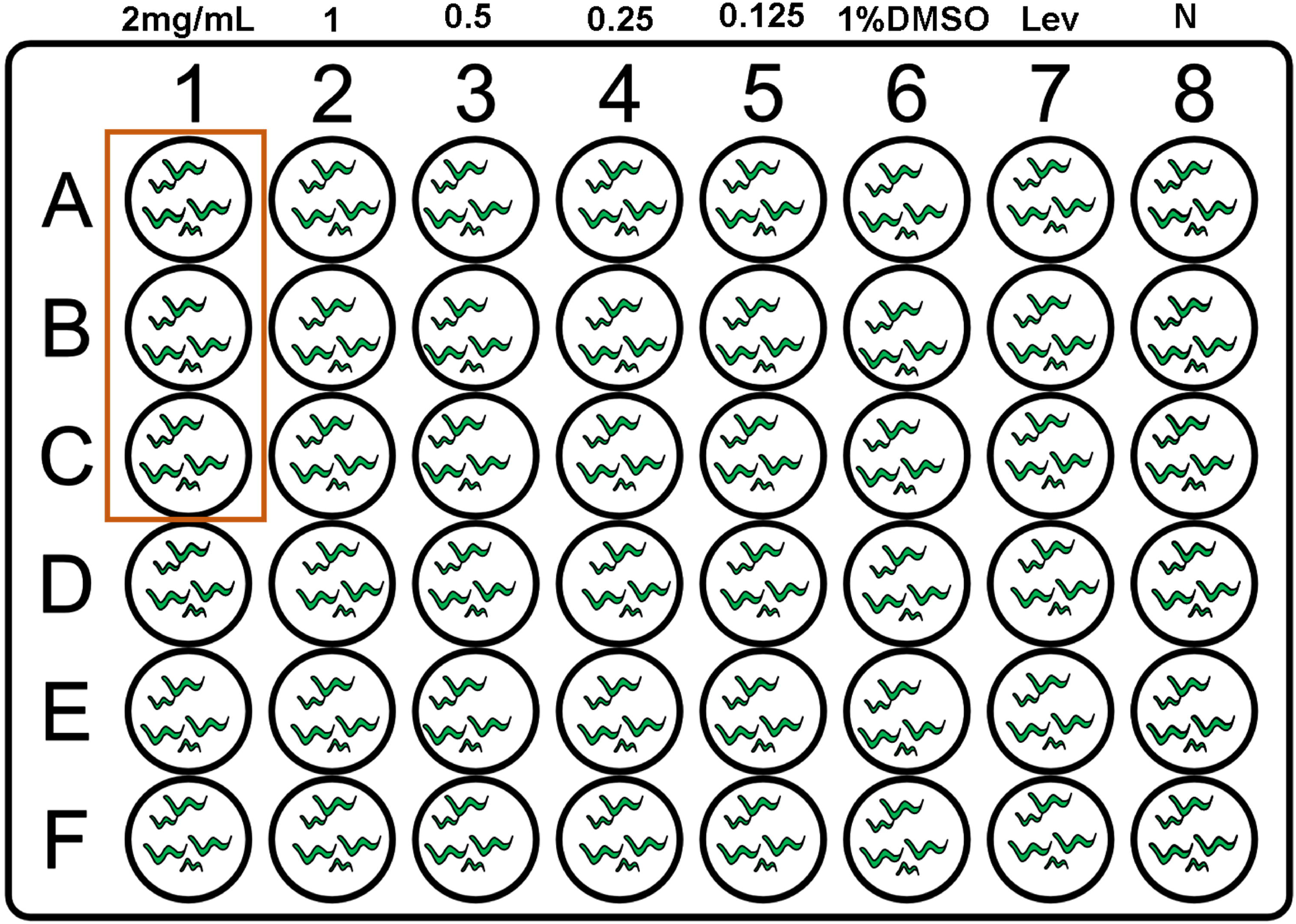
The experimental plot showing the schematic diagram for the evaluation of anthelmintic potency of natural products from medicinal plants at designated concentrations. For every concentration of extracts three replication were taken along with a negative control (N-1% DMSO) and positive control (0.1 mg/mL Levamisole).

**S1 File**. https://doi.org/10.5281/zenodo.17190076

All data and analysis scripts supporting this study are archived and available at https://doi.org/10.5281/zenodo.17190978

